# Cardiolipin increases the peak of reversible traveling H^+^ fronts at the membrane surface

**DOI:** 10.64898/2026.08.15.744977

**Authors:** Nabil-Bey Baroudi, Sergei G. Kruglik, Philippe Lopez, Sinan Haliyo, Stéphane Genet

**Affiliations:** Sorbonne Universite - Universite Paris/Est Creteil, 75005 Paris, France; Laboratoire Jean Perrin, 75005 Paris, France; Institut de Systematique, Evolution, Biodiversite (ISYEB), Sorbonne Universite, CNRS, Museum National d Histoire Naturelle, EPHE, Universite des Antilles, Paris, France; Institut Des Systemes Intelligents et de Robotique, Sorbonne Universite, Paris, France

**Keywords:** cardiolipin, protonation fronts, full field pH imaging

## Abstract

Cardiolipin (CL) is a phospholipid found in the inner mitochondrial membrane (IMM) where it increases the efficiency of ATP regeneration. We have investigated the hypothesis that this increase may result in part from CL concentrating H^+^ at the IMM surface through electrostatic interactions as the CL polar head is a dianion at physiological pH. To this aim, we compared the concentrations and movements of H^+^ at the surface of giant planar phosphatidylcholine (PC) membranes and 20% CL enriched PC membranes by recording their surface pH with the membrane-grafted pH probe fluorescein DHPE. CL enrichment of the membranes increased their surface H^+^ activity by a ∼4 factor. Moreover, we observed non-gaussian spatial H^+^ concentration profiles with distance from a point H^+^ source with both PC and CL membranes suggesting that both lipids also induce interactions between probe molecules. A whole bath pH variation revealed that these interactions allow the traveling of reversible acidification fronts with constant speed over the membrane between high and low pH states. A reaction-diffusion model of these observations suggests that membranes support these fronts through a mechanism of autocatalytic (de)protonation of the membrane surface. In mitochondria, these fronts would result in transitions between high and low pH states, the low one having a larger H^+^ concentration in CL-enriched regions of the IMM. Such an increase at the inner leaflet of the IMM may increase efficiency of the respiratory chain whereas the increase at the outer leaflet may boost the ATP synthase rate.

## INTRODUCTION

Cardiolipin (CL) is a cone-shaped phospholipid whose polar head has two phosphatidic acid moieties and which is found in the inner mitochondrial membrane (IMM). CL has a positive effect on the ratio of ATP regenerated per dioxygen consumed (1). CL concentrates into IMM regions of largest curvature (2). This lipid is thus found in regions of the inner leaflet of the IMM were complexes of the respiratory chain use the free energy liberated by the oxidation of NADH and other organic and mineral compounds to expel H^+^ from the matrix into the inter membrane space. This H^+^ flux results into a H^+^ electrochemical potential gradient across the IMM. CL also concentrates in regions of the outer leaflet of the IMM where the ATP-synthases use the free energy dissipated by the reentry of H^+^ into the matrix to synthesize ATP according to the classical theory of Mitchell (3). Haines (4) has reviewed these features and others suggesting that CL may be crucial for mitochondrial production of ATP via oxidative phosphorylation (OXPHOS) and this hypothesis has found support in studies showing that deficits in CL expression is key in several metabolic diseases including Barth’s syndrome (5, 6). CL is actually not mandatory for the OXPHOS working. It is rather thought to increase the OXPHOS efficiency by, at least, two mechanisms. Firstly, CL stabilizes assemblies of respiratory complexes and is required for the association of the ADP/ATP carrier with respiratory complexes (5) which may increase both the efficiency of electron flow in the respiratory chain and the ADP/ATP exchange. Secondly, Haines and Dencher (7) have proposed that CL may function as a H^+^ trap by condensing H^+^ close to its negatively charged polar head. This would increase the supply of protons to the ATP-synthases. This attraction may also increase the efficiency of the respiratory chain in establishing the inner transmembrane H^+^ gradient. This view is strengthened by the finding that the CL headgroup is fully ionized as a dianion within the physiological pH range and therefore should exert a strong electrostatic attraction on neighboring H^+^ (8). Whilst this view focus on the migration part of H^+^ movements, the question arises as to whether CL may also increase the H^+^ diffusion near the membrane surface since the proteins involved into the mitochondrial production of ATP are spatially distant (9). For instance, Klotzsch et al. (10) have challenged the view that the so-called uncoupling protein 4 may act as a direct uncoupler of oxidative phosphorylation by showing that this protein is spatially separated from both proton pumps and ATP synthases by micrometric distances. This conclusion relies on previous results of Serowy et al. (11) suggesting that, while H^+^ exhibit diffusion coefficients along the surface of lipid bilayers nearly as large as in free solutions, structural H^+^ diffusion (underlain by the Grotthuss mechanism, see eg (12)) is physiologically important only for distances not exceeding 10 nm. However, Weichselbaum et al. (13) have recently provided evidence for an entropic trap of H^+^ at the surface of lipid bilayers that may explain the high affinity of H^+^ for membrane-water interfaces. This trap would channel the movements of highly mobile H^+^ along the membrane surface over distances as large as ∼100 µm and thereby couple spatially distant sites of H^+^ release and consumption in the IMM, in contradiction with the conclusions of Serowy et al. (11). More recently, Weichelsbaum et al. (14) have reported that the charge of the polar head of artificial phospholipidic membranes has no major influence of the H^+^ lateral diffusion coefficient at the surface of these membranes and suggested that the surface charge alone is a poor regulator of proton traffic along the membrane surface.

These diverging results prompted us to undertake a compared investigation of H^+^ movements near the surface of artificial planar phospholipidic membranes either lacking CL or enriched in CL. To this aim, we delivered brief pulses of a UV laser to release H^+^ from NPE-caged-proton close to the surface of membranes build with various lipidic compositions. In order to resolve the spatial distribution of the membrane response to H^+^ pulses, we monitored the H^+^ concentration at various distances from the H^+^ source (0.5 µm resolution) with the membrane grafted, fluorescent pH probe fluorescein-DHPE. Our experiments show that the H^+^ diffusion coefficient close to 20% enriched CL membranes is 35% larger than close to pure PC membranes. This increase appears too small to resolve the problem of coupling OXPHOS proteins by H^+^ fluxes set by the spatial separation of these proteins. Nevertheless, our experiments also revealed that the probe pKa is shifted upward by 0.6 units in 20% CL containing membranes with respect to pure PC membranes. This shift corresponds to a 4-times increase in the membrane H^+^ surface concentration by CL. Moreover, our multi-site superficial pH recordings evidenced non-gaussian spatial H^+^ concentration profiles along the surface of both pure PC and PC-CL membranes. This finding implicates that diffusion alone cannot explain our observed spatiotemporal H^+^ concentration dynamics and thereby suggested that phospholipids were modifying characteristics of the probe (de)protonation reaction. We investigated this hypothesis by monitoring membrane surface pH changings after modifying the pH of the solution bathing the membranes (bulk) as this removed diffusional limits of the focal H^+^ pulses experiments. Decreasing the bath pH triggered acidification fronts that started at points on the membrane edge, travelled at speeds of hundreds of µm per second and eventually acidified the entire membrane surface. This acidification could be fully reversed by increasing the pH back to its initial value which occurred through alkalinization fronts. Thin scratches to the membranes slowed down the fronts but did not prevent their overall propagation suggesting an active propagation mechanism. Simulations of a reaction-diffusion model based on the classical KPP-Fisher equation (15) proves capable to reproduce features of acidification fronts suggesting that phospholipids mediate an autocatalytic protonation of the pH probe. Reversibility of the fronts was straightforwardly reproduced by multiplying the reaction term in the KPP-Fisher equation by a sign term deciding the reaction direction according to the difference between the probe’s pKa and the surface pH.

Our experiments confirm the hypothesis that CL increases the H^+^ diffusion coefficient at their surface (1). However, this increase for the lipidic composition of the IMM (16, 17) would only raise the characteristic length of superficial H^+^ diffusion to 15 nm, a value remaining two orders of magnitude smaller than the maximum distance between OXPHOS proteins. Thus, our experiments conclude that the increase of the coupling between these proteins by CL does not rely on an effect of CL on H^+^ diffusion. Our results suggest that CL rather increases the mitochondrial ATP production through a ∼4 times increase in the number of protons available for both H^+^ transport from the matrix into the intermembrane space by the respiratory chain and at the mouth of ATP-synthases. Most importantly, our experimental setup, specifically designed to resolve the spatial profile of H^+^ concentration dynamics, has allowed us to unravel genuine fronts of acidification that were hypothesized in a previous study (18). These fronts appear as actively propagated all-or-none state transitions that reversibly switch the whole surface or our artificial membranes between high and low H^+^ concentration states. In mitochondria cristae, the spatially uniform low pH state theoretically would overcome limitations imposed by H^+^ diffusion to the coupling of OXPHOS proteins implicated into the mitochondrial ATP synthesis.

## METHODS

### Experiments

#### Experimental setup

Our experimental setup was built around a Nikon microscope and two light sources (Fig.1A): an argon lamp which was used to excite the membrane grafted, fluorescent pH probe fluorescein-DHPE (F-DHPE from Fisher) and a UV laser (10 mW, 405 nm) which was used to photoactivate NPE (1-(2-Nitrophenylethyl sulfate sodium salt from Biotechne), a photosensitizer dissolved into the membranes bathing solutions, which releases a proton upon photolysis. Both light entries passed through the same objective and had the same focal plane. All salts, lipid extracts and reactants were purchased from Sigma. See ***Supp1***.

**Fig 1.**
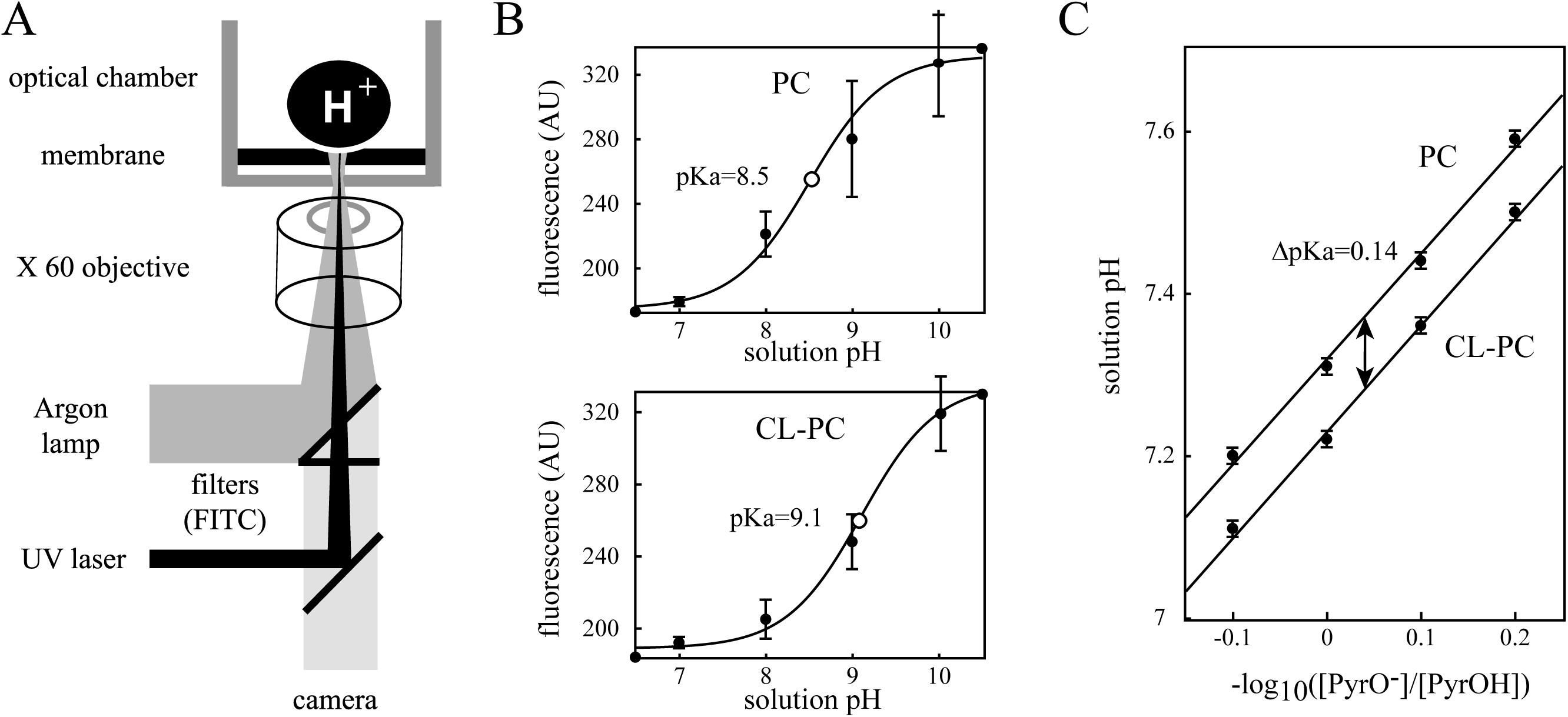
CL acidifies the membrane surface. **A. Experimental set-up**: an inverted microscope with two light entries was used to (i) record the fluorescence of pH-dependent dyes and (ii) to photorelease H^+^ at the surface of planar phospholipidic membranes**. B. Fluorescence of the pH probe fluorescein-DHPE (F-DHPE) grafted to artificial membranes as a function of bath pH.** Solid curves illustrate the results of a nonlinear regression of data against a chemical model of the probe fluorescence (see text). The inflexion point in each curve (open black circles) corresponds to the probe pKa. The largest pKa in CL-enriched membranes as compared to pure PC membranes evidences an acidification of the membrane surface by CL. **C**. **Fluorescence of the pH sensitive dye pyranine (Pyr) inside small unilamellar vesicles versus the decimal logarithm of the fraction of ionized probe molecules,** [PyrO ^−^] / [PyrOH]. Straight lines represent the best regression line against data. Notice that CL decreases the probe pKa, implicating that the probe sees as less acidic solution.

#### Solutions

Three aqueous solutions were used to form giant phospholipidic membranes (19) and to record their superficial pH: *A* (vesicles formation: 55 mM KCl, 50 mM NaCl, 1.1 mM CaCl_2_ (19–21), 250 µM pH buffer (Hepes and Imidazol / Trizma with 3:1 ratio), pH=7.2), *B* (vesicles fusion into SLB) 60 mM KCl, 60 mM NaCl, 3 to 6 mM CaCl_2_, 180µM pH buffer, pH=5.5) and *C* (pH recording at low buffer concentration: 50 KCl, 50 NaCl, 2 mM CaCl_2_, 105µM pH buffer and pH adjusted to 9.2).

#### Vesicular suspension

Small unilamellar vesicles (SUV) were formed by the extrusion method (22–25) or by sonication. From a 10 mg/mL lipid stock in chloroform, 1 mg of freshly nitrogen dried lipid was mixed in a glass vial with 4 mL of aqueous solution A containing 0.5mM pyranine and vortexed for 20 s. The multilamellar vesicles obtained at this stage were then passed through an extruder (Polar Adventi, polyethylene membranes with 100 nm pores) and two Hamilton syringes. We performed 10 passes (26). Final lipidic concentration was 250 µg/mL.

#### Planar phospholipidic membranes supported on glass

A microscope glass slide was plasma treated for 70 s and then immediately mounted in a 2 mL O-ring optical chamber (1.2 cm diameter). A 40 µL droplet of pure water was deposited firstly on the slide surface and 300 µL of a 100 to 250 µg/mL of vesicle suspension (27–30) were then added to the top of the droplet. The addition of 1.2 mL of solution *B* (acidic) induced the fusion of vesicles (27, 31) into a membrane that stuck to the glass slide (29, 32). The optical chamber was then covered (to avoid dust deposit), stored at 37 °C for 20 min and finally rinsed ten times with experimental solution *C* (33) as gently as possible (to avoid destruction of the membrane).

#### Membrane characterization

We used fluorescence recovery after photobleaching (FRAP, see SM#1 (34)), recovery after mechanical scratch (see SM#2 (34)) to verify the correct formation of supported lipid bilayers.

See ***Supp2*** and ***Video3***.

#### Measurements of the pH and H^+^ diffusion coefficient close to the membrane surface

NPE-caged-proton (dissolved at 100 to 250 µM in solution C) was photoexcited by the beam of a UV Laser (10 mW, 405 nm, 50-300 ms duration pulses) to release protons into the bathing solution close to the membrane. A camera (Andor 50 fps) was used to record the pH variation on the membrane surface as acidification decrease the fluorescein signal and fluorescein was chemically grafted in the lipidic composition. Temperature was maintained at 18°C in all experiments.

### Modeling

#### Mathematical modeling of H^+^ concentration dynamics in experiments

Our artificial membranes and their bathing solutions represent a 3d experimental system. However, this system has a symmetry axis going through the point of nucleation of fronts and parallel to the direction of fronts propagation in Experiment#2 (see Results). It follows that the modelization of our experimental system can be reduced to a 2d mathematical problem. Let *Ox* denote the axis of fronts propogation. Let *Oy* denote an axis perpendicular to *Ox* with elevation *y* = 0 corresponding to the bottom of the bathing solution. Our model is made of three connected subsystems and their corresponding state variables

*(M)*: membrane surface (containing the *pH* probe), *d* (fraction of deprotoned probe molecules)

*(S)*: solvation layer of the membrane surface, 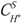 (proton concentration in *S*)

*(B)*: solution bathing the membrane 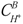 (proton concentration in *B*).

The time and space dynamics of these state variables obey the following set of partial differential equations

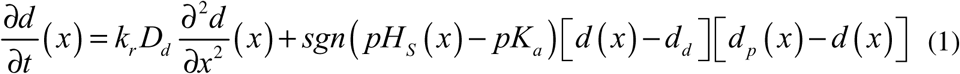

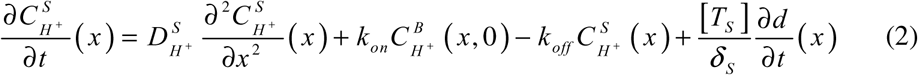

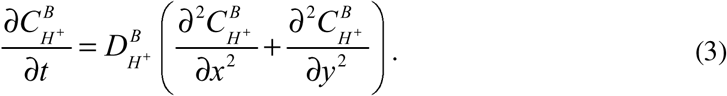

into which 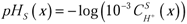 denotes the *pH* at location *x* in the solvation layer and *sgn* stands for the *Sign* function. Symbols *d_d_* and *d_p_* denote two (pH dependent) *d* values corresponding to the probe being mostly in its deprotonated and protonated forms respectively. Derivation of the PDE system and its bounday conditions are detailed in Supplementary Materials.

## RESULTS

### Experiment #1: CL acidifies the membrane surface

To investigate whether CL affects the steady distribution of H^+^ in solutions bathing phospholipidic membranes, we measured the fluorescence of the pH probe fluorescein-DHPE (F-DHPE) inserted in artificial giant membranes with different phospholipidic compositions. Fig1A schematizes our experimental setup (see methods). Fig.1B compares measurements of the F-DHPE fluorescence in pure PC membranes and in 20% CL-enriched PC membranes as a function of the bath pH. Fluorescence intensity, *I*, data in Fig.1B were fitted by function 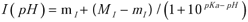 with nonlinear regression using the Levenberg-Marquardt algorithm to minimize differences between experimental and theoretical data (35). The m *_I_* and *M _I_* symbols respectively represent the minimum and maximum fluorescence intensities while the probe pKa corresponds to the pH value were the *I* (*pH*) function admits an inflexion point. The standard errors on pKa estimations were computed according to 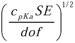 into which *c_pKa_* is the diagonal coefficient corresponding to the pKa in the covariance matrix, *SE* is the total standard error and *dof* is the number of degrees of freedom (number of data points minus number of model parameters). This analysis yiedeld *pKa* = 8.5 ± 0.1 in pure PC membranes and *pKa* = 9.1 ± 0.1 in CL-PC membranes. The pKa difference between PC and CL-PC membranes is highly signifcant since even the 0.1% confidence intervals on pKa estimations do not overlap. This finding strongly supports the conclusion that CL increases the probe pKa. This increase implicates that, for a given pH value, the fluorescence intensity of the probe is smaller in CL-enriched membranes than in pure PC membranes. Given that this intensity is proportional to the *d* = [*FO* ^−^] / [*T_s_*] fraction (13), it follows that *d* must decrease when the local pH decreases and thus that CL acidifies the membrane surface. We obtained insights on the mechanism underlying this acidification by measuring the bulk pH inside small unilamellar vesicles (SUV, 50 nm radius) with the fluorescent dye pyranine (see Methods) as exposed below. Fig.1C shows that straight lines fit well the relation between the solution pH and the decimal logarithm of the the fraction of ionized probe molecules, [*PyrO* ^−^] / [*PyrOH*] in both PC and CL-PC SUV (Fig.1C). Using the Henderson-Hasselbach equation (36), one deduces from data in Fig.1C that CL decreases the pyrnanine pKa by 0.14 unit compared to PC (t-test, p<1/1000). CL therefore apparantly alkalinizes the solution inside SUV (<50 nm) whereas it acidifies the solution close to planar membranes (<2 nm). The electric double layer theory provides an explaination for these opposite results as follows. PC is the major phospholipid in biological membranes (16). Its polar head is made of a choline residue and of a phosphate group which are expected to form a zwiterrion at physiological pH from their pKas. However, experiments have shown that the PC polar head has actually a net -1 charge at physiological pH (37). Owing to thermal agitation, cations (including H^+^) do not fully neutralize the membrane surface charge. The membrane surface is instead surrounded by a cloud of counter-ions which form a diffuse electric double layer (38). Cations in this layer have a larger concentration than in the bulk solution whereas anions exhibit the oppposite behavior. In titration experiment of F-DHPE (Fig.1b), the larger negative surface charge density of CL-PC membranes with respect to pure PC membranes readily explains the F-DHPE pKa increase by a H^+^ enrichment close to the membrane surface through electrostatic attraction. Our ratiometric observation of a CL-induced pKa shift of pyranine in the opposite direction is not contradictory since pyranine is not grafted to the membrane but is a solute in the internal SUV solution. Thus, hydrogen bonding of PyrOH molecules with CL can be predicted to reduce the PyrOH concentration into SUV whereas electrostatic repelling of PyrO^-^ molecules from the membrane surface is expected to increase the bulk concentration of this form of the probe. These two effects may add at increasing [*FO* ^−^] / [*FOH*] ratio and thereby explain the the observed pKa downward shift.

### Experiment #2: Focal release of H^+^ near the membrane surface triggers acidification disks whose radius growth is limited by diffusion

After having shown that CL affects the steady distribution of H^+^ near the membrane surface, we investigated the putative effects of CL on the H^+^ movements at the membrane surface. We addressed this question by measuring the dynamics of the superficial pH in response to brief pulses of H^+^. The superficial pH was again probed using the fluorescence of F-DHPE while membranes were bathed with solutions containing 100 to 250 µM of NPE-caged-proton. This chemical compound has affinity to the membrane and releases H^+^ upon UV photoexcitation at 405nm. The laser beam was focalized at the membrane surface. Fig.2A-B illustrate the typical time evolution of the probe fluorescence (*n*>30) during (A, starting from green) and after the pulse (B, ending by green). The H^+^ liberation began by decreasing locally the probe fluorescence, witnessing a local acidification of the membrane surface (Fig.2A). This acidification had increased by 17% and spread to a ∼15 µm distance from the illumination zone 100 ms after the pulse onset. Later on, during the pulse, the membrane acidification kept increasing (29% after 200 ms and 45% at the pulse end) and spreading (55 µm after 200 ms and 65 µm at the 300 ms pulse end). Fig.2C compares the early (40 to 140 ms from the pulse onset) time evolution of the squared disk radius, *r* ^2^ (µm^2^), sampled at 20 ms intervals (50 fps) in PC and CL-PC membranes (*n*=6 for each data point). Data in both membrane types were well fitted by a straight line (Pearson’s coefficient of determination *R* ^2^ > 99% and p-value of the coefficient test <0.1). We found the linear relations *r* ^2^ = 19*t* − 202 in PC membranes and *r* ^2^ = 26*t* − 340 in CL-PC membranes (*r* in µm and *t* in ms). Such linear relations were expected if diffusion limits the disk expansion (see appendix) and, if so, the slope of these relation reads as 4 *D*, into which *D* (µm^2^s^-1^) is the diffusion coefficient. A regression slope test (p<0.1) showed that the slope in CL-PC membranes is larger than in pure PC ones and thus that CL increases the H^+^ diffusion coefficient near the membrane surface (Fig.2D). This coefficient increases from 4.7 ± 0.5 10^5^µm^2^s^-1^ with pure PC membranes to 6.5 ± 1 10^5^µm^2^s^-1^ with CL-PC ones. Thus, the 20% enrichment in CL of PC membranes increased *D* by 35%.

**Fig 2.**
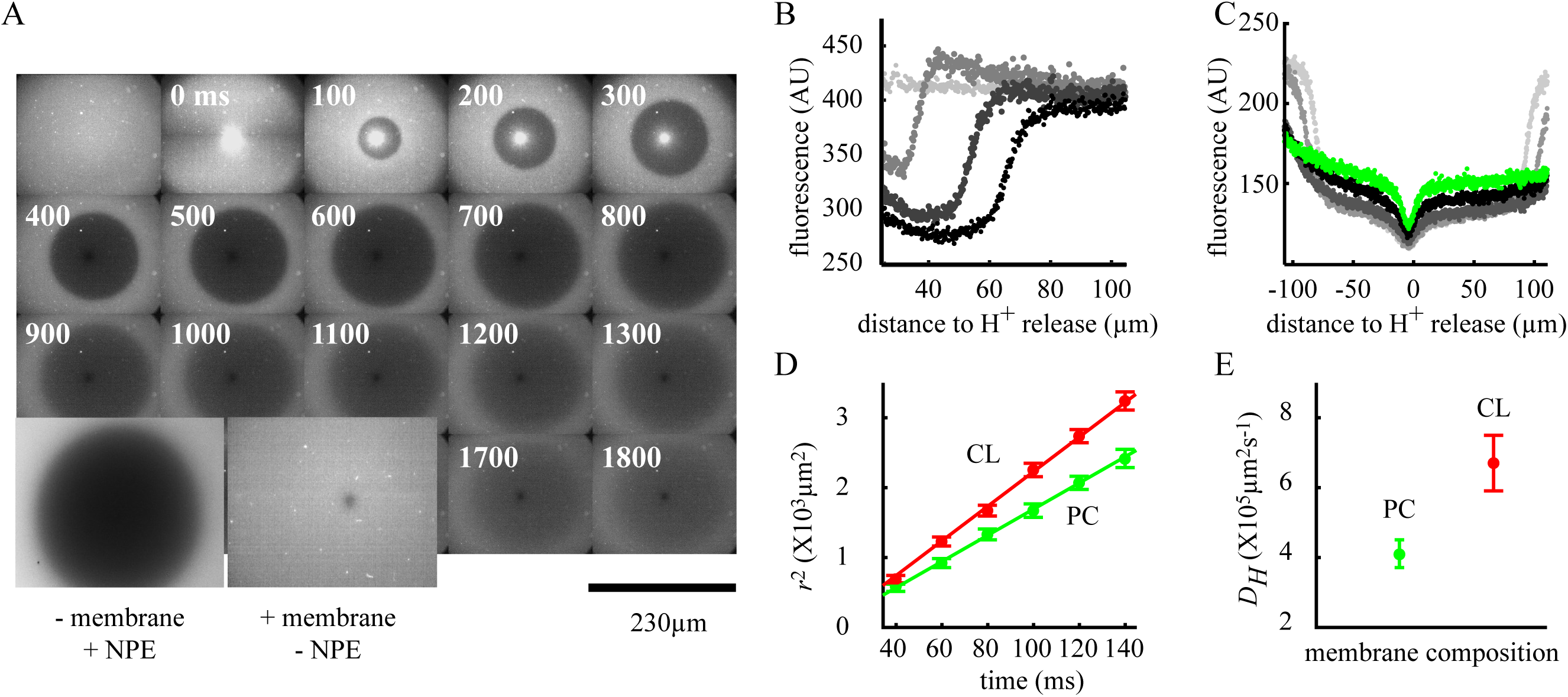
Response of the membrane surface pH to a local release of H^+^ near the membrane. **A.** Snapshots of F-DHPE fluorescence as seen from above the membranes during and following a 300ms H^+^ pulse triggered by photoexcitation of NPE-caged-proton. **B**. Corresponding time evolution of the fluorescence profile along a membrane diameter during the pulse; t=0 (light grey, pulse onset), 100 ms (medium grey), 200 ms (grey) and 300 ms (black, end of pulse). **C**. Snapshots of the fluorescence profiles relaxation from the pulse ending up to 2.5 s at different times after the pulse: 400 ms (light gray), 500 ms (grey), 1 s (medium grey) and 2.5 s (green). **D**. Time evolution of the square radius, r^2^, of acidification disks; CL-PC (red) and PC (green). **E**. Diffusion coefficient computed from the slopes of curves in panel **C**; same color code as in **D**.

The slope of the *r* ^2^*vs t* relationship began to decrease from 160-180 ms, even for 300 to 500 ms pulses. It vanished between 300 to 400 ms, at which the acidification disk reached a maximum diameter of 110-120 µm (Fig.2A). These features resulted from a progressive exhaustion of the released H^+^ as shown by the less wide acidification disks (70-80 µm) triggered by shorter (100 ms) duration pulses (not illustrated). From this moment, the acidification disks began to fade away. Fig.2B shows that the F-DHPE fluorescence profile was nearly flat over the acidification disk diameter during the relaxation phase and thus that the membrane superficial pH was nearly uniform over the disk surface. However, this profile should have had a gaussian shape (see appendix) is diffusion was the unique physicochemical mechanism determining the time and space dynamics of the acidification disks. Thus, it appeared that if diffusion is limiting the early disk expansion, this mechanism was insufficient to explain features of the acidification disk relaxation. We reasoned that membrane/water interface modifies dynamics of the (de)protonation reaction of the probe, 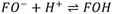, in order to explain our observations during the disk relaxation phase.

### Experiment #3: bulk pH changes trigger reversible waves of membrane surface pH changes in PC and CL-enriched membranes

The above results prompted us to perform a third experiment designed to characterize the mechanism of the putative phospholipid-induced modification of the (de)protonation reaction. This experiment consisted in measuring the dynamics of the superficial pH response to pH changes of the whole bathing solution (instead of focal H^+^ pulses) to remove the diffusion limitation evidenced by experiment #2. In a first step, we deposited a small drop (30-60 µl) of a concentrated HCl solution (50 mM with 10 mM buffer) on the surface of the solution bathing membranes. The solution was then gently homogenized by pipetting for 2-3 s and the evolution of the membrane surface pH was monitored with the F-DHPE fluorescence. After delays of 17.0± 2.4 s (*n*=3, PC) / 18.3 ±1.4 s (*n*=3, CL-PC), an acidification of the membrane surface occurred spontaneously at a point located on the field edge. The difference between PC and CL-PC was not significative (*p*=0.35). From this point, a front of acidification propagated radially at 395 ±58 µms^-1^ (*n*=8, PC membranes) / 283 ±35 µms^-1^ (*n*=3, CL-PC membranes) until the entire membrane surface became acidified. The top of Fig.3A illustrates a representative example of such an acidification front by depicting the fluorescence evolution into a 240x240 µm box at 125 ms time intervals. The black arrow indicates the direction of front propagation while the figure evidences the constant travelling speed of the front. In a second step, once the whole membrane surface had been acidified, a small drop of concentrated NaOH was added to the bath. After delays similar to the onset of the acidification front (19.3 ±1.4 in PC / 17 ±2.4 in CL-PC, no significative difference: *n*=3, *p*=0.2), an alkalinization of the membrane surface occurred at a point on the membrane edge or right under the acidic flow. This local alkalinization turned into an alkalinization front that travelled over the membrane surface at 447 ±48 µms^-1^ (*n*=8, PC membranes) / 430±126 µms^-1^ (*n*=3, CL-PC membranes) (Fig.3A bottom, the white arrow indicates the travelling direction). The speeds of acidification and alkalinization did not differ either in PC (t-test, *p*=0.8, *n*=8) or in CL-PC membranes (*p*=0.3, *n*=3). Moreover, the speed of acidification (alkalinization) did not differ significantly between PC and CL-PC membranes (*p*=0.8 for acidification and *p*=0.7 for alkalinization). Like acidification fronts, alkalinization fronts eventually extended over the whole membrane surface. The nucleation points of both acidification and alkalinization fronts occurred apparently at random as we were unable to identify any geometrical pattern into the set of nucleation points (*n*=10). Fig.3Ba shows that acidification fronts are active events as front propagation were able to overcome a mechanical scratch inflicted to the membrane (top) or a natural defect in the membrane structure (bottom). The propagation of such fronts has been shown to rest on interactions between diffusion and nonlinear reactions in numerous biological processes (39, 40). However, the protonation of the probe, 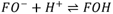 is a first order reaction and has thus a linear reaction rate. We investigated its putative modification by phospholipids into a nonlinear reaction by measuring the fluorescence transfer between two probes, a method that differs from FRET(Förster (or Fluorescence) Resonance Energy Transfer (41)) as it has several seconds duration. Fluorescein (pKa = 6.5 in solution) and Texas Red (pKa <4), were simultaneously grafted to our membranes. The membrane surface was illuminated by blue / UV light (which excites fluorescein) while recording the fluorescence of both dyes (Fig.3C left illustrates the example of a CL-PC membrane). The large intensity used resulted into an extinction of the F-DHPE fluorescence (red trace) by photobleaching. A surge of Texas Red fluorescence (green) coincidated with this extinction, suggesting that the fluorescence lost by F-DHPE was transferred to Texas Red. This transfer can only occur if the distance separating molecules of the donor-acceptor pair is in the range 1-10 nm (42, 43). The right pannel in Fig.3C compares the fluorescence transfer from F-DHPE to Texas Red in pure PC and CL-PC membranes for pH ranging from 7 to 10. The transfer does not exceed 3% over the entire pH range. The transfer is larger in CL-PC membranes over the entire pH range, reaching 14% at pH=8 (and 25% for 5% of each of the probes). This result shows that CL effectively reduces the distance between the dyes in the donor-acceptor pair and enhance transfers. It implicates that, in the presence of CL, probe molecules are less independent of each other than with other phospholipids. For the (de)protonation of *F-DHPE*, this would implicate that the reaction becomes nonlinear and that its rate depends quadratically on concentrations if the reaction follows its stoechiometry. We use this premise in the following section to build a model suggesting that acidification fronts rely on an autocatalycic protonation mechanism.

**Fig 3.**
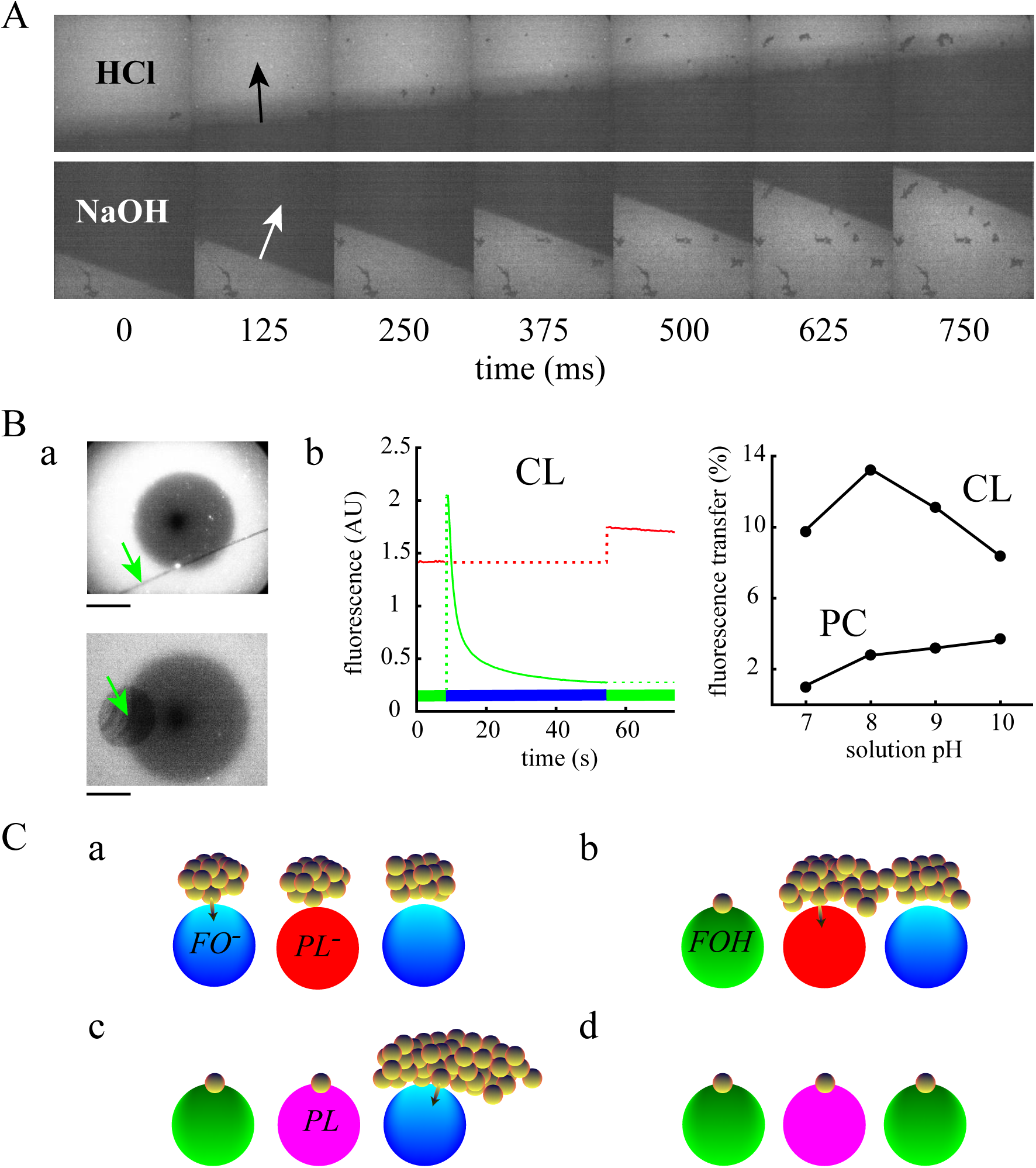
Bulk pH changes triggers reversible fronts of acidification of the membrane surface. **A. Top:** A drop of HCl (40 µl, 50 mM) was deposited on top of the solution bathing a CL-PC membrane and the solution was homogenized. The figure displays the probe fluorescence in grey scale (black: pH=6.5 and white: pH=9.5). After a delay of ∼25 s, a nucleus of acidification occurred at a point of the membrane edge. From then, the nucleus extended into a wave of acidification that ultimately acidified the whole membrane from pH=9.2 to pH=6.7. **Bottom**: a drop of NaOH (40 µl, 50 mM) was subsequently added to the bath solution and the solution was homogenized again. A front of alkalinization spontaneously started from the membrane edge and invaded the entire membrane. Arrows indicate the direction of fronts propagation. **B**. a. **Top**. Fluorescence after a cut (20 µm width black line pointed by green arrow) was inflicted to the membrane. The acidification proves capable to propagate beyond the cut and is only delayed. **Bottom**. A natural defect in the membrane (green arow) neither prevents acidification to extend over the membrane. **b.** Left: fluorescence transfer between fluoresceine (green) and Texas Red (red) grafted to the membranes. Illumination of a CL enriched membrane by blue/ UV light (30-60 s duration indicated by blue horizontal bar) photobleached F-DHPE. The intensity lost by fluorescein is transferred to Texas Red. Right: the fluorescence transfer between dyes is larger in CL-PC than in PC membrane over the pH range 7-10. **D**. Proposed mechanism for reversible waves illustrated in **A**. See text.

**Fig 4.**
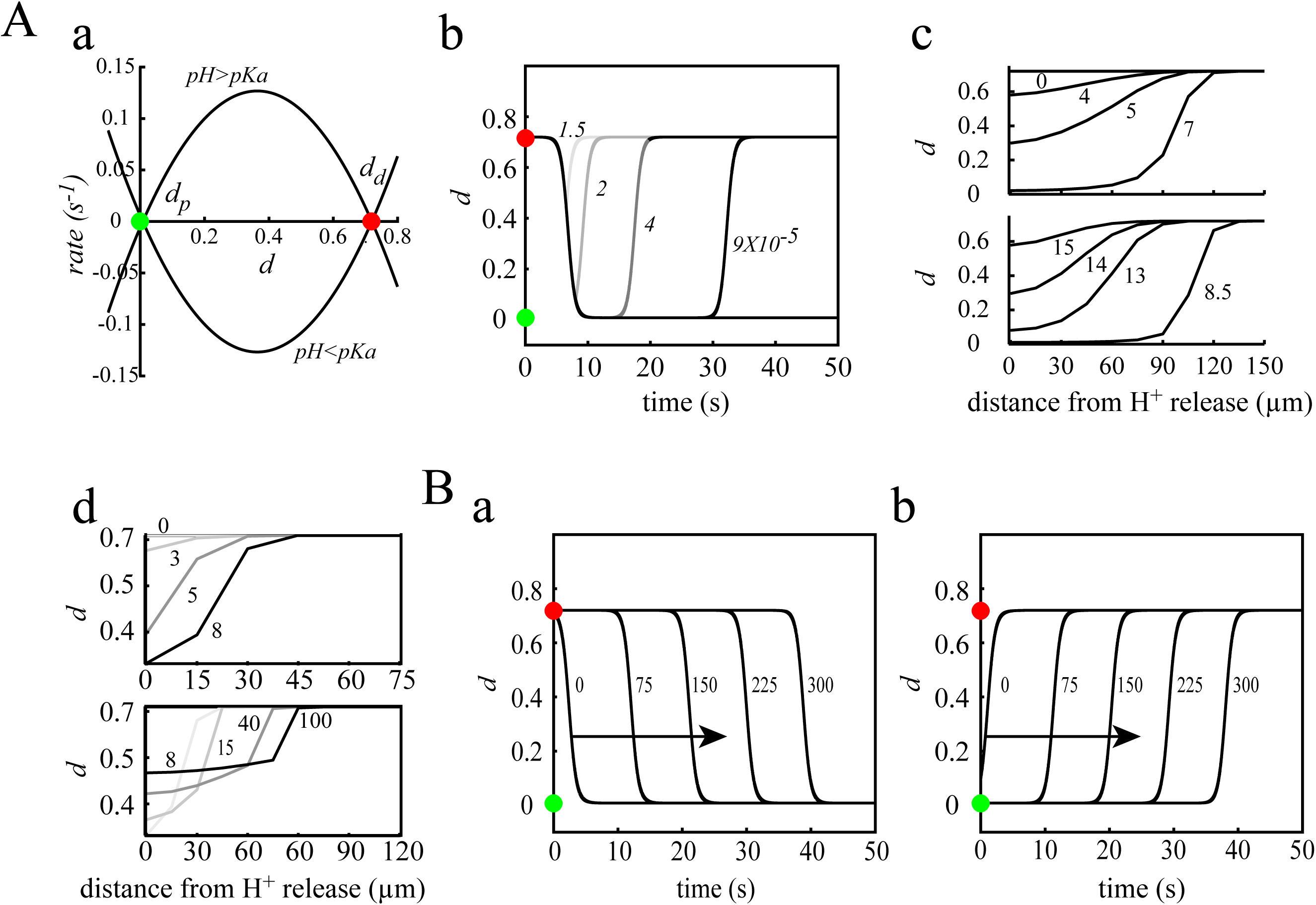
A model of autocatalytic protonation of membrane surface explains results of experiments #2 and #3. **A. Experiment #2. a.** Rate of the autocatalytic reaction (see *Eq. (3)*) as a function of the fraction, d, of deprotoned probe molecules. The reaction has two equilibrium states: deprotonated (d_d_ ≈ 0.72, red dot) and protonated (d_p_ ≈ 7.9 × 10^−3^, green dot). Membrane surface pH changes induce a stability exchange: d_d_ is at pH > pKa whearas d _p_ is stable at pH < pKa. **b and c.** Simulations of experiment #2. **b**. Dynamics of d in response to 300 ms duration pulses of H^+^ delivered at the center of the spatial domain just above the solvation layer. Labels indicate the magnitudes of pulses (Ms^-1^). **c**. Snapshots of the d profile at different times from the start of a 300 ms H^+^ pulse. D. Same as c with d depending on the H^+^ concentration. Top: time expansion of acidification disks. Bottom: relaxation of acidification disks. **B. Experiment #3. a**. Acidification waves. The middle of the bath at its top was locally acidified (pH=7.3 into a 15 µm side square) and the H^+^ concentration was allowed to homogenize by diffusion for 50 s. A 1% perturbation of the d value at edge of the spatial domain triggered a wave of acidification of the membrane surface. Arrow indicates the direction of wave propagation. **b**. Alkalinization waves. After the membrane was entirely switched to d _p_, the model was given 50 s to stabilize its state variable and a 15X15 µm rectangle was alkalinized at pH=13 at top of the middle of the bath. The H^+^ concentration was allowed to homogenize by diffusion for 50 s. A 10% perturbation of d at the left edge of the spatial domain triggered a wave of alkalization turning back the entire membrane surface to the initial d_d_ state of protonation.

Fig.3C illustrates a putative mechanism of interactions between CL and the fluorescence probe *F-DHPE* through electrostatic interactions that may explain the apparent negatively charged phospholipid-induced facilitation of the FO^-^ form protonation by the FOH form (i.e. autocatalysis). The figure displays the theoretical sequence of events leading a pair of FO^-^ molecules to transform into a pair of FOH molecules in response to a pH decrease from 9.2 to 6.7. At the initial pH (9.2), both probe molecules and the polar head of phospholipids are mostly in deprotonated forms (see Fig.1B showing that the probe pKa is 9.1 whereas the CL headgroup is fully ionized as a dianion at pH >5 (8)). Fig.3C displays excess H^+^ (with respect to their bulk concentration) as small yellow balls corresponding to the situation after the drop of concentrated HCL has been deposited. Red balls represent FO^-^ anions whereas magenta balls represent FOH molecules. The situation depicted in Fig. 2Ca corresponds to thermal equilibrium. There exists an electric field **E** whose component *E* _⊥_ normal to the membrane surface is directed toward the membrane and attracts H^+^. The corresponding migration flux of H^+^ (Ohm’s law) is equilibrated by a diffusion flux in the opposite direction. The mean **E** component, *E*_||_, parallel to the membrane surface must be zero to grant equilibrium. The local H^+^ enrichment increases as the pH is decreased to 6.7. This effect increases the probability for a H^+^ to collide an FO^-^ molecule and protonate it through thermal agitation (Fig.3Ca). Neutralization of the FO^-^ charge unravels an *E*_||_ component of the field directed toward the PL^-^ molecule. Canceling of the electrostatic field of the FO^-^ molecule leaves unequilibrated the diffusion flux directed toward the bulk solution and this flux tends to move H^+^ away that were attracted by the FO^-^ molecule. However, diffusion being a slow mechanism, these H^+^ are rather attracted toward the PL^-^ molecule by. This effect further increases the H^+^ concentration near the PL^-^ molecule and to the neighboring FO^-^ molecule at its right. This increases the probability for the PL^-^ molecule to be protonated and, after this protonation, the vicinity of the FO^-^ molecule becomes even more enriched in H^+^ than before the protonation of the left FO^-^ molecule (compare Fig.3Cb and c). In turn, the larger H^+^ concentration increases the probability for the right FO^-^ molecule to be protonated and both probe molecules become protonated in the final state (Fig.3Cd). In summary, the H^+^ enrichment near the 2^nd^ FO^-^ molecule increases the protonation of this molecule, resulting into an apparent autocatalysis of FOH molecules.

### A model of H^+^ concentration dynamics support an autocatalycic protonation mechanism

On the one hand, experiment #1 shows that, in response to a focal H^+^ release, the surface of our artificial membranes develops acidification disks whose expansion is limited by H^+^ diffusion. The disks growth also involves a reaction component as disks exhibit a nearly uniform pH over their surface whereas the pH should have a gaussian shape if diffusion only was at work. On the other hand, experiment #2 reveals that, in condition of uniform bath pH (which removes the limit imposed by diffusion to disks growth in experiment #1), acidification spontaneously occurs at the membrane edge and expanding disks of acidification turn into traveling fronts of acidification. In addition, these fronts appear reversible as alkalinization waves spontaneously start from the membrane edge after the bath pH is increased back to 9.2. Finally, thin scratches of the membranes do not prevent waves propagation and instead result in propagation delays. All of these dynamical properties are reminiscent of those of the KPP-Fisher reaction-diffusion. Actually, the sole qualitative difference between solutions of the KPP-Fisher equations and our experimental results is the reversibility of wave propagation with our membranes whereas state transitions in the KPP-Fisher equation are irreversible. This prompted us to investigate whether the KPP-Fisher equation modified to allow reversible state transitions could reproduce, at least qualitatively, the results of our experiments. Our model is exposed in detail in the Method section. Briefly, the model is 2d owing to the central symmetry of our membranes and it comprises a solvation layer (S) adjacent to the membrane (M) whom surface contains molecules of a protonable fluorescent probe. The solvation layer exchanges H^+^ with a bathing solution (B) according to reactions which are first-order with respect to H^+^ concentrations. The association rate of H^+^ to (S) is *k_on_* (s^-1^) whereas *k_off_* (s^-1^) is the dissociation rate constant. According to previous studies, *k_on_* > *k_off_* so that (S) is acidified with respect to (B). Table 1 lists the parameters of the model, their significance and their default values.

**Table 1.** Parameter values for simulations of the model.

| Symbol (units) | Meaning | Value with reference |
| --- | --- | --- |
| $pH_B$ | pH of the bathing solution in experiment 2 | 9.2 |
| $pKa$ | $pKa$ of the fluorescent probe (fluorescein) | 9 in CL membranes (this study)<br>8.5 in PC membranes (this study) |
| $D_{H^+}^B$ ($m^2s^{-1}$) | Proton diffusion coefficient in bath | $8 \times 10^{-9}$ (53 and this study) |
| $D_{H^+}^S$ ($m^2s^{-1}$) | Proton diffusion coefficient in the solvation layer | $6 \times 10^{-9}$ (derived from experiments of this study) |
| $D_d$ ($m^2s^{-1}$) | Lateral diffusion coefficient of probe molecules over the membrane surface | $1 \times 10^{-11}$-$1 \times 10^{-12}$ (54) and FRAP in *Supp2*). |
| $\delta_s$ (m) | Thickness of the membrane solvation layer | $10^{-8}$ (55) |
| $k_{on}$ ($s^{-1}$) | Rate constant of $H^+$ ‘trapping’ in the solvation layer | 2.3 (13) |
| $k_{off}$ ($s^{-1}$) | Rate constant of $H^+$ release from the solvation layer | 0.5 (13) |
| $k_r$ ($s^{-1}$) | Rate of the autocatalysis reaction | 1 to 10s (adjustable) |
| $[T_s]$ ($Mm^{-2}$) | Membrane surface concentration of probe molecules | $3 \times 10^{-10}$ (estimated) |

In order to dissect out the mechanisms underlying our experimental observations, we performed initial simulations of the model assuming that factor *d_p_* in our modified KPP-Fisher equation (Eq.3) is constant and does not depend on the pH in (S). We simulated experiment #1 by firstly letting the model relax to equilibrium (i.e. until its state variables had reached constant values). State variables are 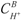 (Mm^-3^), the proton concentration in the bath, 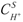 (Mm^-3^), the H^+^ concentration in the solvation layer, and *d* (dimensionless), the fraction of protonated probe molecules. With the bath pH set to 9.2, 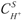 stabilized at 1.62 × 10^−6^ *Mm* ^−3^ corresponding to a superficial pH=8.79 and the fraction of deprotoned molecules was *d_d_*=0.72. At this pH value, which is smaller than the probe pKa, the deprotoned form is dominant and the *d_d_* state is stable. We then investigated the membrane response to brief (300ms duration) pulses of H^+^ delivered just above (S) at the center of the spatial domain in order to mimick the release of H^+^ by NPE photolysis in experiment #1 (Fig.4Ab). A 1.5 ×10^−5^ *Ms* ^−1^ pulse only triggered a brief, tringular shaped acidification of the membrane surface as witnessed by a transient *d* decrease. Increasing the pulse magnitude to 2 ×10^−5^ *Ms* ^−1^ increased both duration and magnitude of the acidification. Increasing the magnitude to 4 ×10^−5^ *Ms* ^−1^ resulted in a saturation of the acidification with *d* decreasing to the value 7.9 ×10^−3^ corresponding to the *d_p_* state. The membrane remained in this state for ∼5 s after which the acidification faded away and the membrane surface reconvered the *d_d_* state. A further increase in the pulse magnitude lengthened the period over which the membrane stayed in the *d _p_* state (∼20s with 9 ×10^-5^ *Ms*^-1^ in Fig.Aab). Subsequent increases in the pulse magnitude evidenced an exponential dependence of the time spent by the model in the *d _p_* state on the pulse magnitude (not shown). This time became infinite with very large pulses and corresponded to an amount of released H^+^ sufficient to keep *pH_S_* > *pKa* even after the (slow) dissociation of H^+^ from probe molecules was completed (as demonstrated by the flat *d* profile at 7.9 ×10^−3^ at the end of simulations). Finally, we simulated the time evolution of the *d* profile in response to a H^+^ pulse so as to compare the model previsions with the results of experiment #1 in Fig.2A-B. Fig.4Ac displays snaphots of the *d* profile taken at differents instants from the begnning of a 300 ms pulse of H^+^. Top of the figure illustrates the expansion phase of acidifications disks while the bottom of the figure illustrates the relaxation dynamics of the disks. Labels on the horizontal axis indicate the distance from the region of H*^+^* release. The flat profile at time *t* = 0 corresponds to a membrane surface over which protonation of the probe has an identical value *d_d_* = 0.72. At time *t* = 4 s, the center of the membrane has begun to acidify, as shown by the *d* decrease close to the H^+^ liberation site. One second later, the acidification has increased and has begun to spread from the membrane center owing to the H^+^ lateral diffusion. At time *t* = 7*s*, *d* has decreased to a minimum value which is nearly identical over distances from the membrane center ranging from 0 to ∼60 µm. These results reproduce well the experimental expansion of acidification disks displayed on Fig.2A. At the end of the expansion phase (time *t* = 8.5*s*), the disk reaches a maximum radius of ∼110 µm. This value is in close agreement with our experimental observations. From this moment, the acidification disk began to fade away through two concomitant processes (compare snapshots at times *t* = 13,14 and 15 s. Instead of further increasing its diameter, the disk shrunk through the effect of lateral H^+^ diffusion as the slow deprotonation of the probe increased back *d* in the central region of the membrane where *d* exhibits its minimum value. The unique qualitative difference between the disk relaxation in the model and in experiment #1 is that *d* begins to increase back while the disk radius keeps increasing in experiments whereas in the model the radius begins to decrease before *d* has begun to increase back. This difference did not originate from the parameter values used for the simulations as we obtained the same results in a broad range of values for the diffusion coefficients (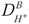 and 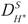) and the rate constants (*k_on_* and *k_off_*). However, recall that results illustrated in Fig.4Ac were obtained assuming that *d_p_*is constant whereas it should actually depend on pH_s_ (see Eq.3bis). We tested whether the *d_p_* dependence on pH_s_ could account for the different *d* dynamics in experiments and in simulations by replicating the simulation illustrated in Fig.4Ac using Eq.3bis for *d_p_*. Fig.4ad illustrates the time evolution of the *d* profile in response to a 10^-4^Ms^-1^ amplitude and 300 ms duration H^+^ pulse with variable *d_p_*. Like the simulation with constant *d_p_* (Fig.4Ac), the model with variable *d_p_* reproduces well the *d* dynamics during the early part of growth of acidification disks (Fig.4Ad top). Morevover, the *d_p_*-variable version of the model correctly accounts for the late part of the disk growth: *d* begins to increase on the membrane around the H^+^ source while the disk radius keeps increasing (Fig.4Ad) like observed in experiment #1 (Fig.2B). Thus, our model using a modified KPP-Fisher equation into which the *d_p_* state depends on pH in the solvation layer (S) accounts for the results of experiment #3. This adequation between the model and experiment #3 gives a first support to our hypothesis according to which negatively charged phospholipids underlie an autocatalytic mecanism of protonation of the membrane surface.

We then simulated experiment #2, into which the bath was acidified by depositing an HCl drop by setting 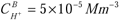 in a 15 µm side square located at the center of the top of the bath. The proton concentration in the model was then left to homogenize by diffusion. Complete homogenization required ∼50 s after which 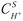 reached a value of 1.63 × 10^−6^ *Mm* ^−3^. This value corresponds to an 8.79 superficial pH, a value laying just below the probe pKa (i.e. 8.8). In the model, the deprotonated state, d_d_ (see Eq.3), becomes unstable at this pH (see Fig.4Aa). Despite this instability, the membrane remained in the d_d_ state over protracted times (tested up to 10 mn). However, a small perturbation (*δ d* = 10^−2^) of *d* at the left boundary of the spatial domain triggered an acidification wave that invaded the entire domain (Fig.4Ba) and eventually switched the whole membrane surface to the protonated *d_p_*state. The acidification wave traveled at a speed of ∼100 µms^-1^ in the model. We then checked whether the model could also reproduce the reverse alkalinization waves observed experimentally. Fistly, the model was left to equilibrate in the *d _p_* state for 10s. Then, the top of the bath was alkalinized into a region of 15 ×105 µm ([H ^+^] = 10 ×10^−10^ *Mm*^−3^). The H^+^ concentration in the bath was then left to equilibrate by diffusion for 1mn (contradiction). The superficial pH reached a value of 8.81. This value laid just above the probe pKa and thereby made the protonated state, *d*_p_, unstable while it allowed the *d_d_* state to recover stability. Although the *d*_p_ state was now unstable, the membrane remained in this state for arbitrarily long times (tested up to 10 mn). However, a small *d* perurbation (*δ d* = 10^−1^) triggered a wave of alkalinization (Fig.4Bb). This wave invaded the entire spatial domain with a ∼10 µms^-1^ speed and switched back the entire membrane to the *d_d_* state. Thus, our model also proves capable to reproduce the main results of experiment #2. This agreement of the model with experiments further supports our hypothesis of an autocatalytic mecanism of membrane surface protonation and suggests that this mechanism is reversible. This article brings experimental evidences that this mecanism could be Red/Ox dependant (fluorescence transfer) and litterature that potential and hydrogen bondings could play a key role in its occurrence.

## CONCLUSION

We intended our fist experiment to unravel putative CL-induced changes of the membrane superficial pH by comparing the titration curves of the protonable residue of the fluorescein-DHPE (*F-DHPE*) pH probe in pure PC and CL-PC membranes. This experiment shows that CL induces a right shift of the probe pKa by 0.6 units. The definitions of the pH and the pKa implicate that a CL-induced pKa increase of 0.6 pH units correspond to a H^+^ enrichment of the membrane surface by a factor ∼4. On the other hand, the theory of the diffuse electric double layer (38, 44, 45) states that the enrichment factor obeys a Maxwell-Boltzmann statistics and reads 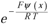, into which *R* (JK^-1^M^-1^) is the gas constant, *F* (CM^-1^) the Faraday constant, *T* (K) the temperature and *ψ* (*x*) (V) the mean electrostatic potential at distance *x* from the membrane surface (with the zero of potential taken at the limit *x* → + ∞). One deduces an electrostatic potential value of *ψ* ≃ − 35mV from the above measured value of the enrichment factor and its theroretical expression. Given that the pKa increase relies upon mobile H^+^, this electrostatic potential value must be compared to the zeta potential (potential at the hydrodynamic plane of shear marking the outer edge of the Stern layer into which ions are tightly bound to the membrane (46)). Sathappa and Alder (8) have measured a nearly twice larger magnitude (-60mV) zeta potential with 20% CL-enriched vesicles. However, Sathappa and Alder bathed their vesicles with solutions having an ionic strength about 6 times smaller than our membrane bathing solution whereas Wnek and Davies (47) report that increasing the ionic strength exert complicated effects on the zeta potential magnitude of colloidal particles depending on the ionic strength range. These effects apparently extend to superficially charged planar liquid-solid interfaces, at least to silica wafer surface, the zeta potential of which decreases as the ionic strength increases (48). It can be estimated from data in (48) that an increase in ionic strength from 20 mM (conditions in (8)) to 118 mM (our conditions) should reduce the zeta potential of silica wafer by ∼15%, i.e. from -60 mV to -51 mV as compared to -35 mV in our experiments. Most importantly, Khalifat et al. (23) report that adding 10% CL to the membrane of PC large unilamellar vesicles (containing only 6mM of salts) decreases their zeta potential from -7 mV to -56.9 mV. Our values cannot match theirs quantitavely given that we used different electrolytic solutions to form our respective vesicles. However, the experimental evidence by Khalifat et al. (23) that CL renders the membrane surface potential more negative supports our conclusion that CL enriches in H^+^ the neighborhood of membranes through electrostatic attraction.

Our H^+^ release pulse experiment shows that a 20% CL enrichment of PC membranes increases the H^+^ diffusion coefficient near the membrane by ∼37% (Fig.2D). It is tempting to bring together this increase and the CL-induced acidification discussed above given that the diffusion coefficient is concentration-dependent in most systems (49). We are unaware of studies investigating this concentration dependence for the diffusion of protons. However, Chakrabarti (50) has reported that the cesium diffusion coefficient monotonically decreases with increasing concentration of CsCl. If the same holds for other cations including H^+^, the CL-induced increase in H^+^ diffusion coefficient cannot be explained by the H^+^ enrichment of the membrane surface in our experiments. The H^+^ diffusion coefficient increase may rather result from an enhancement of the Grotthuss mechanism of H^+^ jumping through the hydrogen bond network of water molecules. Indeed, Scheu et al. (51) have observed that anions (like the CL polar head) are preferentially hydrated with respect to cations. Serowy et al. (11) have estimated that the characteristic length of H^+^ diffusion is up to ∼10 nm. Now, recall that this length scales with 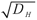 so that the 37% increase in *D _H_* should only moderatly increases the diffusion up to ∼12 nm. Thus, this increase is unlikely to contribute to the coupling between H^+^-dependent proteins of the OXPHOS as these proteins can separated by distances as large as 1 µm.

However, our full field experimental setup allowing to monitor the pH at multiple points from a H^+^ source suggests that diffusion is not the sole the mechanism governing the H^+^ movements at the membrane surface. Indeed, the pH profile with distance from the H^+^ source is not gaussian which implicates that H^+^ traffic close to our membranes is not entirely determined by diffusion (see Appendix). This conclusion is strengthened by the observation that acidification disks triggered by a H^+^ pulse abruptly ended their expansion in a threshold-like fashion. The diffusion equation cannot account for threshold phenomenon as this equation is linear and threshold phenomenon necessarily involve nonlinear reactions. Thus, our results suggested that H^+^ movements at the surface of our membranes involved such a nonlinear reaction. Our experiment of fluorescence transfer between two probes shows that reaction at the membrane surface and reaction in the bulk are coupled by phospholipids, particularly with CL that compensate the bulk protonation by Red/Ox reaction at the surface plane (25% with CL, 6% with pure PC). We attempted to characterize this reaction by changing the pH of the whole bath in order to dissect out the reaction from diffusion. Decreasing the bath pH below the probe pKa triggered travelling fronts of acidification whose propagation was neither impaired by scratching the membranes nor by obstacles to H^+^ movements at the membrane surface constituted by impurities. These features are reminiscent of travelling solutions of the KPP-Fisher equation: constant speed into a homogeneous medium and induction of delays by potential wells as illustrated in Fig. 3B. This prompted to test the hypothesis that negatively charged phospholipids, in particular CL, may underlie an autocatalytic protonation mechanism between probe molecules. We achieved this test by simulating a PDE model of our experiments based on the KPP-Fisher equation for the reaction term. This model proves capable to reproduce features of experimental acidification fronts. It can also reproduce their reversibility through alkalinization fronts triggered by increasing back the bulk pH to its initial value. We obtained front reversibility by multiplying the reaction term in the KPP-Fisher equation by the *ad hoc* factor *sgn*(*pH_S_* (*x*) − *pK_a_*).

Acidification fronts, as well as alkalinization ones, systematically started at a nucleation point on the border of the membrane after latencies in the order of seconds. The localization of this point varied from one experiment to the other thereby suggesting that fronts originated from pH fluctuations at the interface between the membrane and the edge of the optical chamber. This hypothesis is strengthened by the finding that the pH at liquid-solid interfaces can be widely different from the bulk pH (see e.g. (52)). The flow of injected HCl could itself produce nucleation point on the membrane as it produces both potential and fluctuation.

The traveling speed of these waves (larger than 100 µms^-1^) would ensure that CL rich regions of the IMM can respond with <10 ms delays to changings imposed to the OXPHOS by metabolic demands. Taken together, our combined experimental and theoretical results suggest reappraising the OXPHOS regulation by showing (i) that negatively charged polar heads of membrane phospholipids enhance the travelling of reversible fronts of reversible acidification of titratable residues at the membrane surface but are not mandatory and (ii) that CL increase the H^+^ concentration peak of these waves. The former property would overcome H^+^ diffusion limitations imposed by micrometric distances between proton pumps and the ATP synthase molecules while the latter would increase the efficiency of both the respiratory chain and ATP synthesis. All lipids that we did test (PC, PE, PS, DOTAP, DHPE and CL) could produce some waves regardless of their net charge and length but we focused on the effect of CL in this study in regard of OXPHOS metabolism. Understanding those waves could help to treat certain mitochondrial metabolic diseases as protons movements have a central role in mitochondrial metabolism.

## ACKNOWLEDGMENTS

Peter Reinhardt, Thomas Panier and Jean Cognet are gratefully acknowledged for their help in building the experimental setup and contributing to the project design. Fluorescence image acquisition was performed at the IBPS Imaging Facility and authors greatly acknowledge France Lam and Chloé Chaumeton of the Imaging Facility for their expert help. The IBPS Imaging facility is supported by Région-Île-de-France, Sorbonne-University and CNRS. We are also grateful to ENVA veterinary school and to Hôpital Mondor and its IMRB institute.

## AUTHOR CONTRIBUTIONS

N-B.B designed the research and performed experiments. SG designed the model. PL simulated the model. SH contributed to the design of experiments and the data analysis. SK conceived and built the prototype of the experimental setup and helped to characterize signals. All authors participated to the redaction of the paper.

## APPENDIX

### A.#Theoretical evidence for a diffusion limiting process in experiment#1

The 1d diffusion equation describing the time and space evolution of the volume concentration, *C* (Mm^-3^), of a diffusing substance in a 1d spatial domain writes (Crank, 1975)

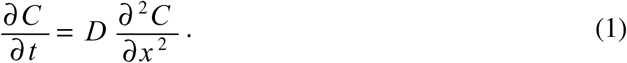

Solution of (1) in an infinite spatial domain with initial conditions 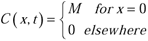 reads (Crank, 1975)

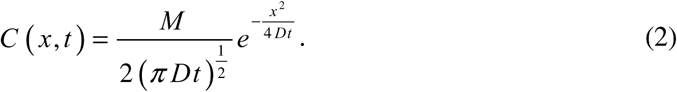

Function (I.2) describes the time and space dynamics of *C* after an amount *M* of the diffusing substance is dropped at time *t* = 0 and location *x* = 0. Let us then consider a Laplace-Gauss function with zero mean and variance *σ* ^2^

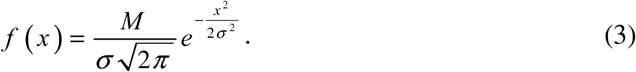

Functions (2) and (3) can be equated by putting *σ* ^2^ = 2 *Dt* into (2) (or *σ* ^2^ = 4 *Dt* in our experiments since they involve a 2d spatial domain). This formal adequation indicates that (*i*) the linear expansion with time of the squared radius, *r*, of acidification disks in Experiment#1 implies that diffusion governs the disks expansion dynamics and (*ii*) that the spatial profile of *C* should be gaussian at any time if diffusion solely underlies these dynamics.

