## Supplementary material for "Cardiolipin increases the peak of reversible traveling H^+^ fronts at the membrane surface": text

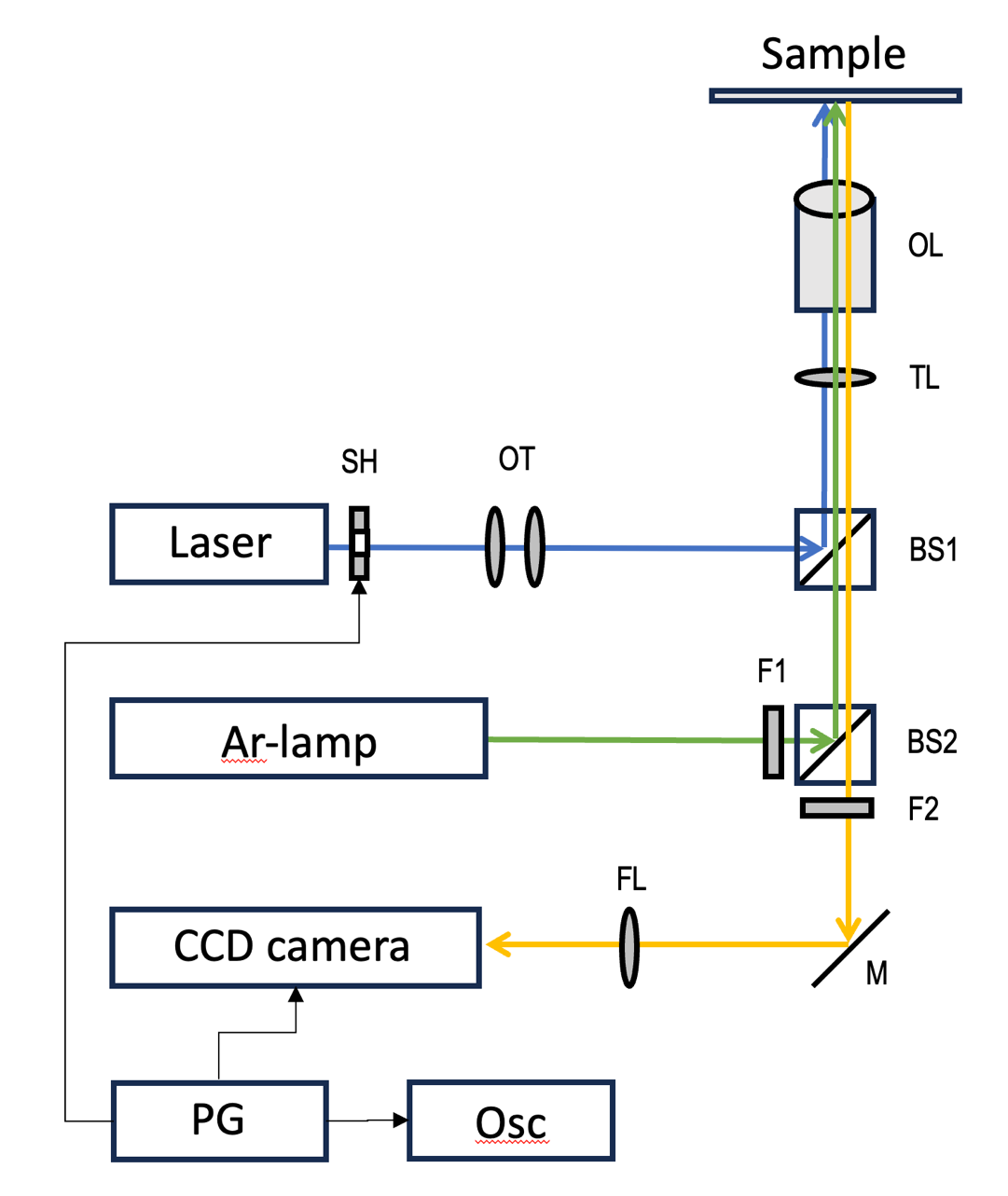

**Supp1.** Schematic presentation of the experimental setup for local proton release: Laser, continuous-wave laser (Coherent) at 405 nm; SH, fast mechanical shutter (Uniblitz), for regulation of temporal length of laser excitation pulse; OT, optical telescope, for regulation of excitation spot size; BS1, dichroic beamsplitter, for laser light reflection and transmission of any longer-wavelengths light; TL, microscope tube lens; OL, objective lens (Nikon 60X, 1.25); Ar-lamp, standard microscope argon lamp in housing with optical collimation scheme; F1, blue-green optical filter for excitation of FL-DHPE fluorescence; BS2, dichroic beamsplitter, for reflection of blue-green excitation light and transmission of green-yellow fluorescence; F2, green-yellow optical fluorescence filter; M, mirror; FL, focusing lens; CCD camera, sensitive CCD detector (Andor); PG, electronic pulse generator (AIM-TTI), Osc, oscilloscope (Tektronix) for pulse monitoring. Laser beam (10 mW, 4 mW measured in the optical chamber at focal point) was used for creation the source of protons that were releases upon photolysis of NPE. The laser beam was focused into the narrow spot of 21 µm radius and with average power density of 300 W/cm2 after focalization on the membrane plan; laser spot size was regulated using the optical telescope OT. The zero-time position and laser pulse duration were regulated using fast mechanical shutter coupled with pulse generator and monitored using an oscilloscope. The continuous weak blue-green light from Argon lamp was used to excite fluorescence of FL-DHPE for monitoring fluorescence intensity during protons propagation. The Argon-lamp illumination spot region of measurements was limited by the size of CDD detector which was used to record the fluorescence signal from FL-DHPE.

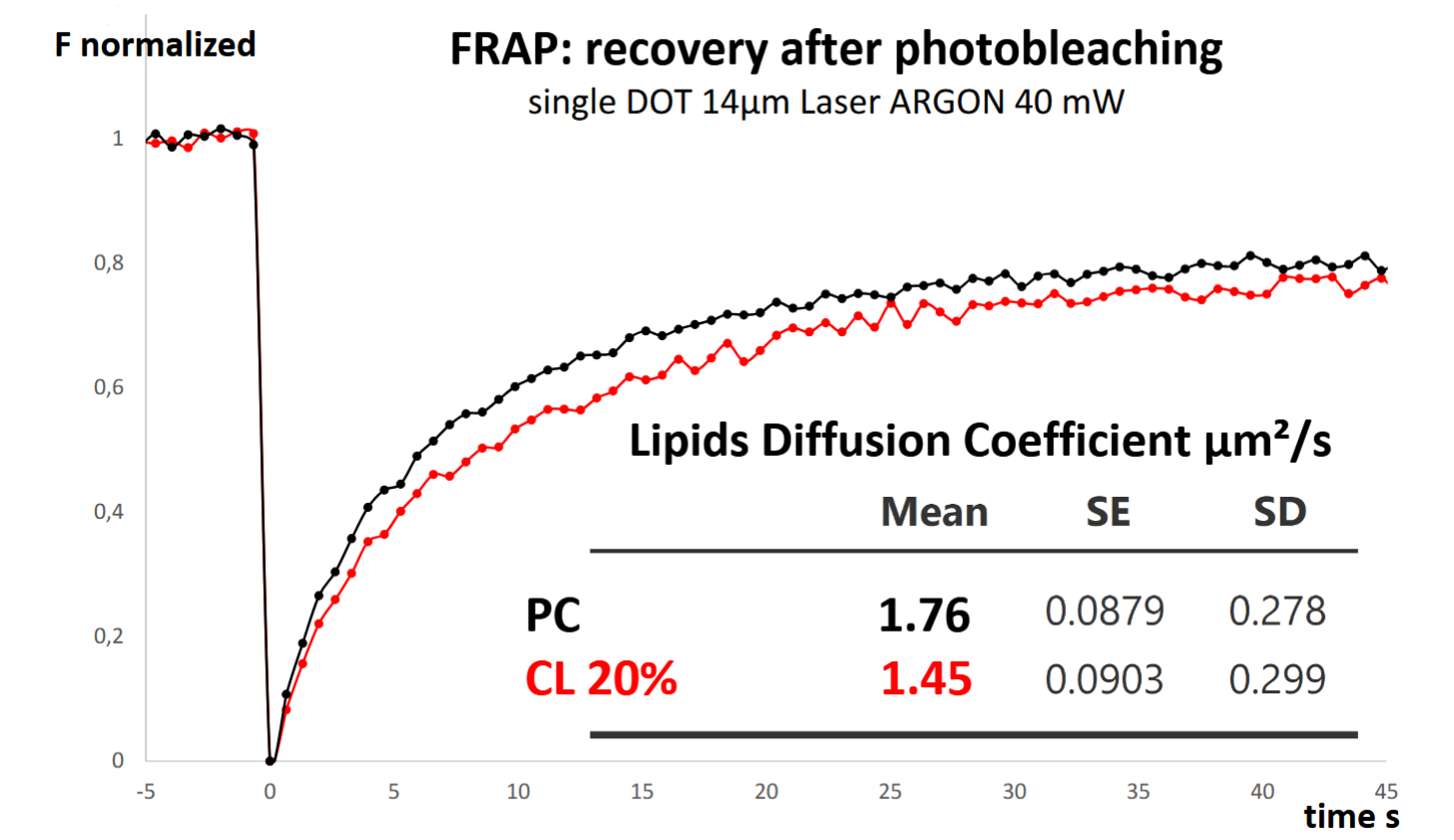

**Supp2:** FRAP shows that lipids within the membranes have 85% mobility and that lipids diffuse slowly within the membrane (compared to protons). The fluorescence signal increases when the membrane is well formed so photobleaching and FRAP are easy to use to check the membrane integrity. n=20 and 20 membranes.

I. **Videos illustrating membrane characterization and reversible fronts of acidification**

***Video #1 (acidification front.avi) :* Example of an acidification front traveling along the membrane surface**

A 40 µL droplet of buffered HCl (0.05 M) is added in the optical chamber, acidifying the whole bathing solution. After 6 seconds from the start of the video, a protonation front (fluorescence extinction) travels over the recording field to its top at 235 µm/s. Video was slowed down 3 times. Notice the curvature of the front. The Leica FRAP microscope was used for its fast linear laser scanning and low photobleaching.

***Video #2 ((reverse) alkalinization front.avi) :* A subsequent drop of NaOH triggers a reverse front of alkalinization**

This is the continuation of video #2. Video starts after the addition of a drop of NaOH to the bathing solution. A deprotonation front (fluorescence recovery) appears 4 seconds after the video beginning. This alkalization front travels at 443 µm/s.

***Video #3 (membrane scratch healing.avi) :* Membrane self-repairs after a mechanical scratch**

This video illustrates the spontaneous repairing of the membrane (when well formed) after a mechanical scratch. This method is complementary to FRAP to assess the membrane integrity and fluidity. Video is accelerated 3 times. Fronts of acidification/alkalinization could cross barriers formed by these scratches before their healing.

**II	Mathematical modeling of H^+^ concentration dynamics in experiments 1#3**

Let
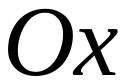
 denote an axis passing through a diameter of the (disk shaped) membrane and
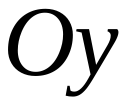
 denote an axis perpendicular to
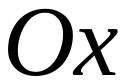
. Elevation
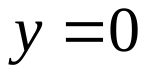
 corresponds to the bottom of the bathing solution (*B*) at which lies the solvation layer (*S*) of the membrane surface (*M*). The membrane surface contains the pH probe fluorescein-DHPE (*F-DHPE*). In solution, fluorescein coexists in protonated (
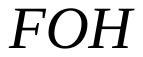
) and deprotonated forms (
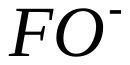
) according to reaction

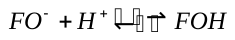
. (1)

Reaction (1) is characterized by equilibrium constant

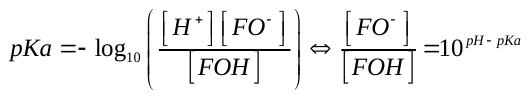
 (2)

with (surface) concentrations of
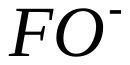
 and
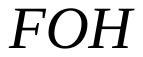
 in Mm^-2^. According to (2), the probe is mostly in its
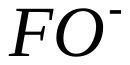
, fluorescent form at
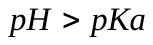
 whereas the non-fluorescent form,
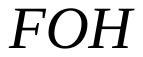
, is dominant at
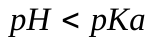
. Let
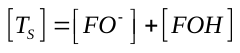
 denote the total probe surface concentration (Mm^-2^). The model assumes that reaction (1) is much faster than the H^+^ exchanges between (S) and (B) and than the H^+^ diffusion in (S). Thus, reaction (1) is considered at equilibrium at any time.

State variables in the model are
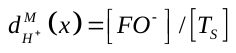
, the molar fraction (dimensionless) of the deprotoned probe surface concentration to the total probe surface concentration,
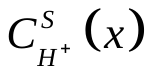
, the H^+^ concentration (Mm^-3^) in the solvation layer and
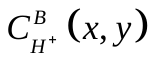
 the H^+^ concentration in the membrane bathing solution (Mm^-3^). We assume that
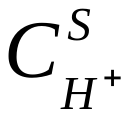
 has a uniform value over the *S* thickness (hence the independence of
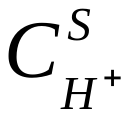
 with respect to *y*) owing to the very small thickness of the solvation layer. The model assumes (*i*) that probe molecules diffuse along the membrane with diffusion coefficient
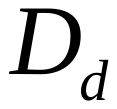
(m^2^s^-1^) and (*ii*) that (de)protonation of the probe obeys the simple quadratic autocatalytic scheme of the KPP-Fisher equation after a modification allowing reversibility of state transitions. According to theses hypotheses, the PDE describing the time-space evolution of *d* reads

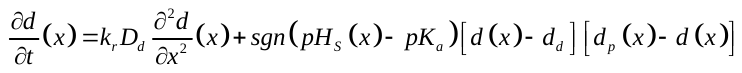
 (3)

into which
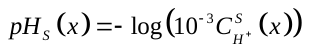
 denotes the *pH* at location *x* in the solvation layer and *sgn* stands for the *Sign* function. Symbols *d_d_* and *d_p_* denote two membrane states into which the probe is respectively in its *FO^-^* and *FOH* forms with

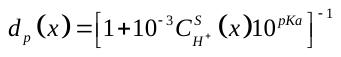
 (3bis)

The deprotoned value of *d*, *d_d_*, was considered constant since our experiments use the same alkakline bathing conditions (pH=9.2). The
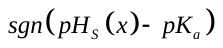
 factor in (3) allows pH_s_ variations to exchange the stability of states *d_d_* and *d_p_*. Parameter
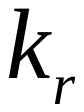
 stands for the rate constant (s^-1^) of the (de)protonation reaction.

**Dynamics of *H*^+^ concentration in the solvation layer and bathing solution**

In order to grant the material conservation law, derivation of PDEs for *H*^+^ concentrations
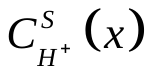
 and
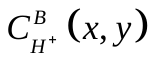
 started from the continuity equation in differential form

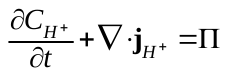
 (4)

into which
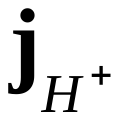
 denote the H^+^ flux density (Mm^-2^s^-1^).
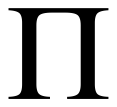
 (Mm^-3^s^-1^) is a source/sink term accounting for chemical reactions involving H^+^ (see below) and
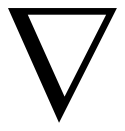
 is the divergence operator. The solvation layer and bathing solution are connected through H^+^ exchanges across their shared boundary at
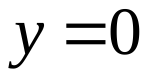
. Therefore, we had first to express H^+^ fluxes through this boundary in order to derive PDEs for H^+^ concentrations. Weichselbaum et al. (13) have shown that H^+^ have a high affinity for the near-membrane water layer. Following their modelization approach, we described the exchange of H^+^ between (S) and the bottom of (B) as the difference between two unidirectional fluxes depending linearly on H^+^ concentrations. The respective densities (Mm^-2^s^-1^) of these fluxes read

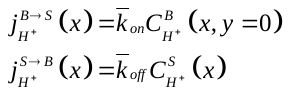
. (5)

Constant
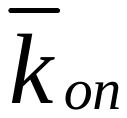
 and
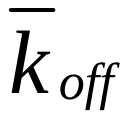
 in (II.3) are pseudo chemical rate constants (ms^-1^).

***Solvation layer***

We derived the PDE for
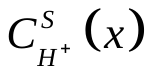
 by assuming that

(i) H^+^ diffuse in (S) along the *x*-axis with diffusion coefficient
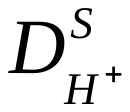
,
(ii) The solvation layer is so thin (ref) that
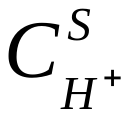
has a uniform value over the layer thickness,
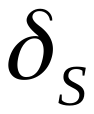
(m), and hence does not depend on *y*,
(iii) the solvation layer exchanges H^+^ with the bathing solution according to fluxes given by (5),
(iv) the solvation layer gains/losses H^+^ through the (de)protonation of the pH probe.

These hypotheses lea to the following PDE

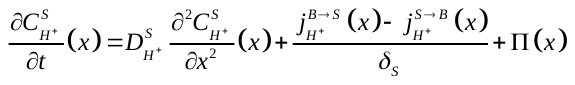
 (6)

According to the above assumptions, the
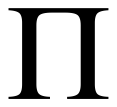
 term reads

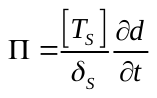
 (7)

into which
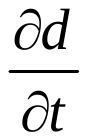
 is given by (3). After inserting (5) and (7) in (6) one finally gets

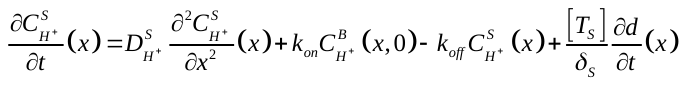
 (8)

into which
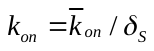
 and
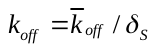
 are rate constants in s^-1^.

***Bathing solution***

Our model assumes that H^+^ diffuse freely into the bathing solution. Accordingly,
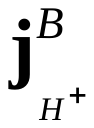
obeys Fick’s first law in (B) and reads

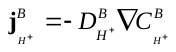
 (9)

into which
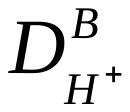
 (m^2^s^-1^) stands for the *H*^+^ diffusion coefficient in water. Substituting (9) for
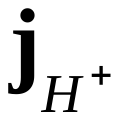
in (4) and taking into account (*i*) that the bathing solution is devoid of chemical reactions in our experiments and (*ii*) our arguments for reducing the model formulation to a 2d problem, the continuity equation leads to the following PDE for
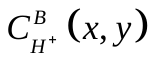

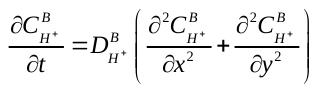
. (10)

PDE (10) holds at any point in (B) with the exception of (B) boundaries. Thus, solving (10) requires formulating boundary conditions. Initial conditions are also required since Eq.s (3,6&10) are infinite dimension Cauchy problems.

**Boundary conditions**

Zero flux boundary condtions were imposed at the left, right and top boundaries of (B) since H^+^ cannot escape the recording chamber

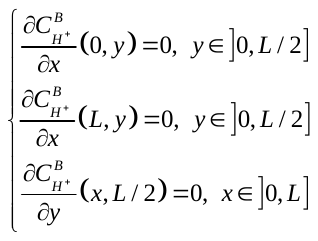
. (11)

We wrote a flux boundary condition at the bottom boundary of (B) (
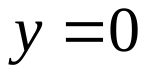
) corresponding to the connection of (B) and (S) by the H^+^ fluxes given by (5)

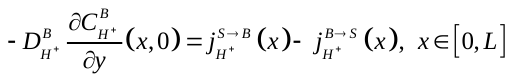
. (12)

**Initial conditions, experimental initial radius and hedge effects on diffusion**

We used the following two sets of initial conditions to simulate our experiments.

Exp#1: acid ‘drop’ provided by NPE photoexcitation

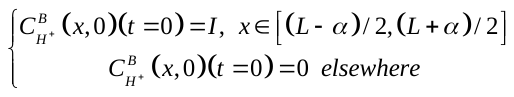
. (13)

Exp#2: uniform bath pH changings

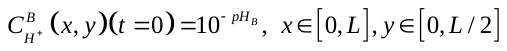
. (14)

The second order spatial derivative terms into the PDE system (3, 8, 10) were discretized with a centered finite difference scheme while boundary conditions were discretized with a first order finite difference approximation. We integrated the resulting system of ODE with the CVODE method implemented into the XPP software. Simulations of the model were achieved into a
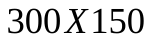
µm rectangle with a mesh size of 15µm (corresponding to the diameter of the region of the bath solution illuminated by the UV laser).
